# WNT-driven immune evasion promotes malignant transformation of *BRAF*-mutant colorectal cancer

**DOI:** 10.64898/2026.01.07.697968

**Authors:** Manuel Mastel, Aitana Guiseris Martinez, Umberto Pozza, Jasmin Meier, Ioannis Chiotakakos, Sandra Jaun, Gabriele Diamante, Nikolaos Georgakopoulos, Philipp Albrecht, Yvonne Petersen, Saskia Reuter, Barbara Schmitt, Michael Günther, Istiffa Nurfauziah, Ian Ghezzi, Kyanna S. Ouyang, Mick Milsom, Jens Puschhof, Nic G. Reitsam, Kim E. Boonekamp, Johannes Betge, Steffen Ormanns, Michael Boutros, Rene Jackstadt

## Abstract

*BRAF*-mutant colorectal cancer (CRC) constitutes a molecularly and clinically distinct subtype with poor prognosis and resistance to standard therapies, representing a major unmet clinical need. Arising from the serrated pathway of colorectal carcinogenesis rather than the classical adenoma-carcinoma sequence, this subgroup remains relatively understudied yet displays a more aggressive disease course. To investigate the progression of serrated CRC, we generated multiple genetically engineered mouse models (GEMMs) of *BRAF*-mutant, microsatellite-stable (MSS) CRC that closely recapitulate human disease. Our findings demonstrate that WNT-pathway activation via *APC*- or *CTNNB1*-mutations, but not *RNF43*-loss, initiates serrated CRC by suppressing immune-mediated tumor surveillance. Mechanistically, WNT-signaling drives distinct alterations in T cell phenotypes within the tumor microenvironment, enabling tumor progression. Together, these data indicate that WNT-signaling mediates immune escape during the malignant transformation of *BRAF*-mutant CRC.

## Introduction

Colorectal cancer (CRC) can arise through distinct molecular and histopathological routes (1–3). In addition to the classical pathway, CRC can develop via a more aggressive serrated route, characterized by precursor lesions with saw-tooth-like histology (1). This serrated route comprises ∼20% of CRCs and represents distinct genetic features, a higher occurrence in the right side of the colon, and precursor lesions with flat morphology, which makes them difficult to detect via colonoscopy, thus representing a clear unmet clinical need (4). Morphologically, serrated precursor lesions can be classified as hyperplastic polyps (HPPs), sessile serrated adenomas/polyps (SSAs/SSPs), and traditional serrated adenomas (TSAs) (5). Initiation events of serrated lesions are activating point mutations in *BRAF* or *KRAS,* leading to enhanced MAPK signaling and increased proliferation (6,7). *BRAF*^V600E^ hotspot mutations are found in ∼10% of CRCs and are associated with SSA precursors and define a clinically distinct subgroup characterized by poor prognosis and low response rates to conventional chemotherapies (8,9). *BRAF*-activated serrated human tumors exhibit characteristic DNA methylation patterns (CpG island methylator phenotype; CIMP) and nuclear β-catenin localization, indicative of tumor-associated activation of the WNT/β-catenin pathway (10,11). Approximately half of the serrated CRCs are associated with microsatellite instability (MSI), which is for example driven by silencing of tumor suppressor genes like *MLH1*; a subtype that shows high response rates to immune checkpoint inhibition (ICI) (12). The remaining 50% of serrated CRCs are microsatellite stable (MSS) and tend to follow a significantly more aggressive course (13).

While estimations predict that 20-30% of CRC arises from serrated precursor lesions, only ∼10% of CRCs actually present with serrated features (14). For better functional understanding of disease progression, genetically engineered mouse models (GEMMs) serve as useful and important tools (15). Mice harboring oncogenic *BRAF* develop widespread serrated hyperplasia that progresses with age to dysplastic serrated adenomas but rarely to metastatic carcinoma. These advanced lesions exhibit enhanced MAPK-activity and activating WNT/β-catenin mutations (6,16). Immunohistochemical analyses of human SSAs show that nuclear β-catenin is found in serrated lesions with neoplastic potential (11). This supports a central role for WNT-signaling during serrated tumor progression. The combination of oncogenic *BRAF* with genes associated with *BRAF*-mutant serrated CRC, such as *CDX2* (17), *CDKN1A* (6,18), *ALK5*/TGFBRI (19), *TP53* (6) and *SMAD4* (20) gives rise to distinct CRC phenotypes, particularly with respect to WNT-signaling activity. Notably, some of the *BRAF*-mutant mouse models develop tumors that lack detectable WNT-activation (1), reflecting heterogeneity in pathway dependence.

Cancer cells influence their surroundings by secreting cytokines, chemokines and growth factors that reprogram immune and stromal cells to adopt pro-tumorigenic phenotypes (22). The composition of factors that are produced by cancer cells may be instructed by cell-intrinsic cues such as their genetics or epigenetics (23,24). Particularly, T cells are central to adaptive anti-tumor immunity, capable of directly recognizing and killing malignant cells (22). Therapeutic strategies targeting *BRAF*-mutant CRC by inhibiting the MAPK-pathway have been proven challenging due to rapid feedback activation and pathway redundancy (25). Combination strategies, such as BRAF inhibition combined with MEK or EGFR inhibition, have shown promise in improving outcomes for patients with *BRAF*-mutant CRC (26). Interestingly, alterations in *RNF43*, a negative regulator of WNT-signaling, showed predictive capacity for response to BRAF/EGFR inhibition (27). Resistance to BRAF/EGFR was found to be driven by ERK-independent SRC-mediated transcriptional reprograming via β-catenin signaling (28). Again, this indicates a central role for WNT-signaling in therapy resistance of *BRAF*-mutant CRC. Recently, therapeutic efficacy has been successfully extended by combining ICI with BRAF/MEK or BRAF/EGFR targeting (29,30). This highlights the potential for combinatorial therapies, including ICIs, and underscores the importance of understanding the tumor immune microenvironment in *BRAF*-mutant CRC.

Limited understanding of the transition from SSA to serrated CRC emphasizes the need for improved mechanistic insight into pathways and processes driving this malignant transformation. Here we generated multiple GEMMs and employed organoid transplantation models to dissect genetic events driving the progression of *BRAF*-mutant CRC. Because the role of WNT-signaling in serrated CRC is poorly defined, we examined recurrent WNT-pathway alterations identified in our GEMMs and patient cohorts. We found that WNT-pathway activation, by *APC* or *CTNNB1* mutations, are required for SSA transformation. Mechanistic characterization showed that WNT-activation drives immune evasion in serrated CRC by impairing T cell phenotypes. Ultimately, this opens opportunities to test novel immunotherapy strategies for WNT-altered MSS CRC.

### Statement of Significance

This study identifies WNT-pathway activation as a critical driver of malignant transformation in *BRAF*-mutant CRC. Using genetically engineered mouse models and single-cell multi-omics, we demonstrate that WNT-signaling impairs T cell phenotypes. This enables tumors to escape immune surveillance, promote disease progression, and reveals a targetable vulnerability in this aggressive CRC subtype.

## Results

### Modeling advanced *Brat*-mutant CRC uncovers WNT-activation

*BRAF*-mutant CRC has previously been associated to poor prognosis. To validate this, we integrated two datasets (TCGA and MSK) (31,32) of human CRC (Figure S1A). Survival analysis confirmed that *BRAF*-mutated CRCs show significantly worse survival compared to *BRAF*-wild type tumors, independently of other alterations or clinical features (Figure S1B). When we compared *KRAS*-mutant versus *BRAF*-mutant CRC, *BRAF*-mutant cases showed significantly lower survival (Figure S1C), underlining the aggressive nature of *BRAF*-mutant CRC. To model this aggressive and metastatic CRC subtype, we intercrossed *Villin1*Cre^ER^ mice with a *Brat*^V600E^ and floxed *Trp53* allele (VBP) (Figure 1A), as there is a high frequency of *TP53* mutations observed in human MSS *BRAF*-mutant CRC (33). Intra-peritoneal injection of tamoxifen and subsequent recombination of floxed alleles generated metastatic tumors (Figure 1B-D; S1D). We previously described that Notch1 activation is a strong driver of metastasis in *Kras*-mutant serrated CRC (7). Therefore, we intercrossed a conditional Notch1 intra cellular domain (N1icd) allele into VBP mice (VBPN). The activation of Notch1-signaling upon Cre-recombination, significantly enhanced metastasis, while no impact on survival or number of tumors was detected (Figure 1B-C; S1E). To further characterize the molecular features of these tumors we analyzed the activation of WNT-signaling by immunohistochemistry (IHC) for β-catenin. In the majority of tumors, we detected an activation of WNT-signaling, manifested by nuclear β-catenin or WNT-target gene expression (*Lgr5*) (Figure 1E-F; S1F-G). To determine whether this WNT-activation is based on genetic alterations, we performed whole exome sequencing (WES) of ten VBPN tumors. Indeed, 40% of the tumors presented genetic alterations in the previously identified CRC driver gene *Ctnnb1*. We found mutations in the phosphorylation sites S33, I35 and T41, located in exon 3. Mutations at those sites prevent degradation and thereby lead to β-catenin stabilization (Figure 1G). Similarly, in human *BRAF*-mutant MSS CRC, we observed that approximately one-third of tumors were nuclear β-catenin positive. Moreover, *CTNNB1* and other WNT-pathway mutations were detected in BRAF-mutant MSS CRC tumors from the TCGA (34) and MSK (35) datasets (Figure 1H-J; S1I). In sum, we have generated highly metastatic GEMMs of *Braf*-mutant CRC that recapitulate the activation of WNT-signaling as detected in human *BRAF*-mutant CRC.

**Figure 1.**
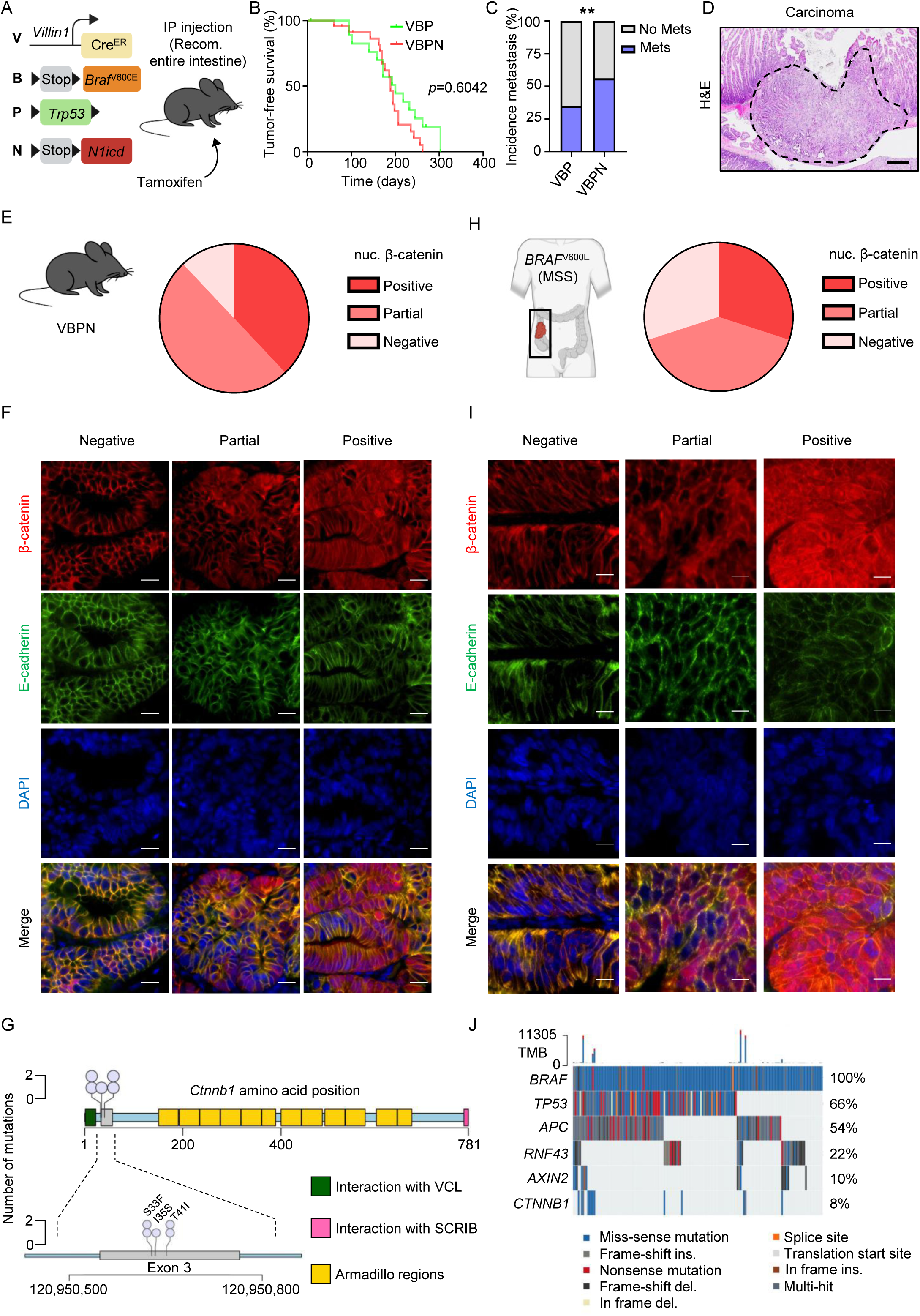
Genetic modeling of *Brat*-mutant colorectal cancer. **A,** Schematic describing the genetics and induction of VBPN mice. **B,** Kaplan-Meier curve of VBP and VBPN mice post induction. Log-rank (Mantel-Cox) test. **C,** Bar-graph showing the incidence of metastasis in VBP and VBPN mice. Statistical significance was assessed by Fisher’s exact test. **D,** Representative hematoxylin and eosin (H&E) image of VBPN carcinoma. Scale bar 100 µm. **E,** Percentage of tumors (VBPN) with nuclear β-catenin expression (n=26). **F,** Immunofluorescence of VBPN tumors. Scale bar 50 µm. **G,** Lollipop plot illustrating genetic alterations identified via WES in tumors from VBPN mice (n = 10). The upper panel displays the full-length protein structure with mutations mapped to their specific amino acid positions. The lower panel provides an expanded schematic of exon 3, detailing the cluster of genomic alterations within this hotspot region. **H,** Percentage of human *BRAF*-mutant MSS tumors with nuclear β-catenin expression (n=10). **I,** Immunofluorescence of human *BRAF* mutant MSS tumors. Scale bar 50 µm. **J,** Oncoprint of human *BRAF*-mutant CRC. Tumor mutational burden (TMB); (n=157).

### WNT-activation drives malignant transformation of serrated CRC

Next, we set out to define the functional role of genetic WNT-pathway alterations in the progression of serrated CRC. To this end, we intercrossed VBPN mice with a floxed allele of *Rnf43*^fl/fl^ (VBPNR), *Ape*^fl/+^ (VBPNA) or an activating deletion of exon 3 in *Ctnnb1*^fl/+^ (VBPNC). To determine whether this additional alteration leads to CRC development, we induced recombination at the end of the distal colon by injection of 4-hydroxy tamoxifen (4-OHT). This resulted in tumor development only at the injection site, with no tumors observed elsewhere in the intestine (Figure 2A-B). This technology enabled us to assess specifically tumor initiation. Longitudinal monitoring showed that only VBPNC and VBPNA mice generated tumors, while VBPN or VBPNR did not within one-year post injection (Figure 2C-D). Tumors arising in VBPNA and VPBNC showed strong nuclear localization of β-catenin (Figure 2E-F). Principal component analysis (PCA) of bulk RNA-seq data of VBPN tumors (IP injection) or VBPNA/VBPNC tumors showed a clear separation of tumors, based on with WNT-activation status (Figure 2G-H). WNT-activation resulted in a subtype switch from the mesenchymal consensus molecular subtype 4 (CMS4) in VBPN to the WNT-high subtypes, CMS2 and CMS3, in VBPNA and VBPNC tumors (Figure 2I; S1H). This shift from the mesenchymal subtype to the WNT-high and metabolic subtypes (8), is in line with the activation of WNT-signaling. Interestingly, activation of WNT-signaling and the subtype switch co-occurred with a down-regulation of Notch1-signaling activity (Figure 2J-K). Again, this agrees with our previous observation that Notch1-signaling is a major driver of CMS4 (7). Furthermore, we identified a strong downregulation of immune response signaling in WNT-high (VBPNA and VBPNC versus VBPN) tumors (Figure 2L). In sum, we provide functional evidence that WNT-pathway activation is a central part of tumor initiation in *Braf*-mutant CRC.

**Figure 2.**
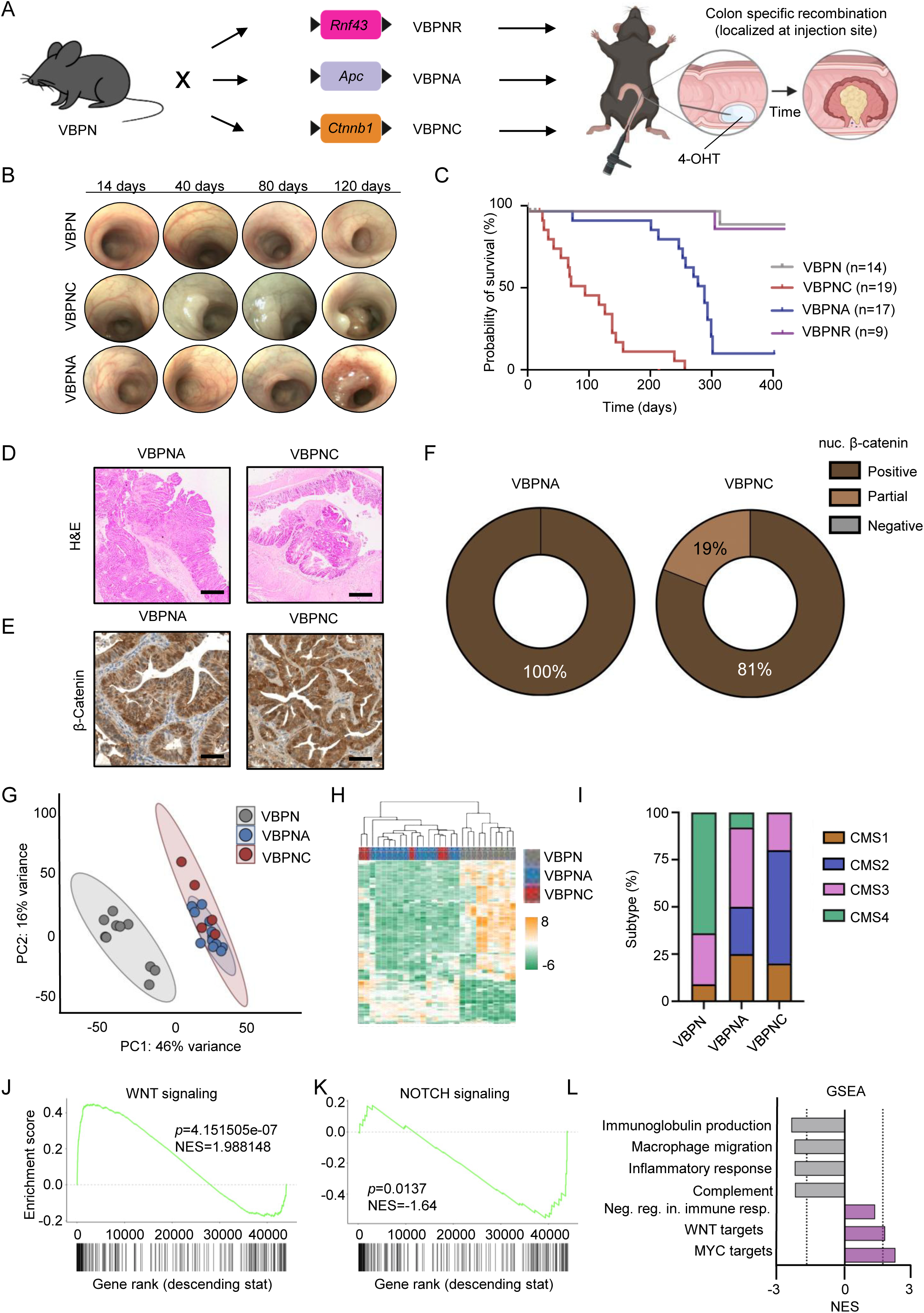
WNT-pathway alterations and tumor initiation in *Brat*-mutant models. **A,** Schematic describing the breeding and induction strategy of VBPN, VBPNR, VBPNA and VBPNC mice. **B,** Representative colonoscopy pictures of indicated genotypes over time. **C,** Kaplan-Meier curve of indicated genotypes post induction as outlined in A. Log-rank (Mantel-Cox) test. **D,** Representative H&E image of VBPNA and VBPNC tumors. Scale bar 400 µm. **E,** Representative β-catenin stain of VBPNA and VBPNC tumors. Scale bar 200 µm. **F,** Percentage of tumors with nuclear β-catenin expression (n=5 VBPNA; n=21 VBPNC). **G,** PCA plot of bulk RNA-seq data across genotypes (n=10 VBPN; n=12 VBPNA; n=5 VBPNC). **H,** Heatmap showing differentially expressed genes across genotypes. **I,** Consensus molecular subtype analysis across the genotypes. **J,** Gene set-enrichment analysis of VBPN versus VBPNA/VBPNC for WNT-pathway activation. **K,** Gene set-enrichment analysis of VBPN versus VBPNA/VBPNC for NOTCH-pathway activation. **L,** Gene set-enrichment analysis of VBPN versus VBPNA/VBPNC for indicated gene sets. Adjusted p values shown in the enrichment plots in J, K and L by applying the Benjamini-Hochberg procedure to control the false discovery rate across all tested gene sets.

### WNT-driven immune evasion *in vivo*

Next, we aimed to functionally investigate the effect of WNT-activation on immune surveillance. To avoid the acquisition of unwanted or undefined driver mutations during tumor progression *in vivo*, we isolated normal colonic epithelial cells from VBPN mice, without prior induction by tamoxifen. After establishing organoids, we transformed the organoids, by adding 4-OHT to the culture medium (Figure 3A). Resulting cultures appeared as spherical organoids that grew also upon withdrawal of EGF from the media (Figure 3A). These recombined cultures were then subjected to gene editing to produce isogenic organoid lines. To create an *Apc* knockout, we utilized conventional CRISPR/Cas9 gene editing with a previously validated sgRNA targeting *Apc* (36). Base editing was used to create the oncogenic S33F mutation in *Ctnnb1* (37) (Figure S2A-C). We isolated two individual clones of each sgRNA accounting for potential clonal effects (Figure 3A). RNA-seq of the resulting cultures revealed strong up-regulation of canonical WNT-targets and negative regulation of immune-regulatory processes (Figure 3B-G, S2D-E). We confirmed activation of the WNT-pathway by increased expression of the WNT-target *Lgr5* (Figure S2F) and increased TOP-GFP reporter activation upon *Apc* and *Ctnnb1* mutation (Figure S2G-H).

**Figure 3.**
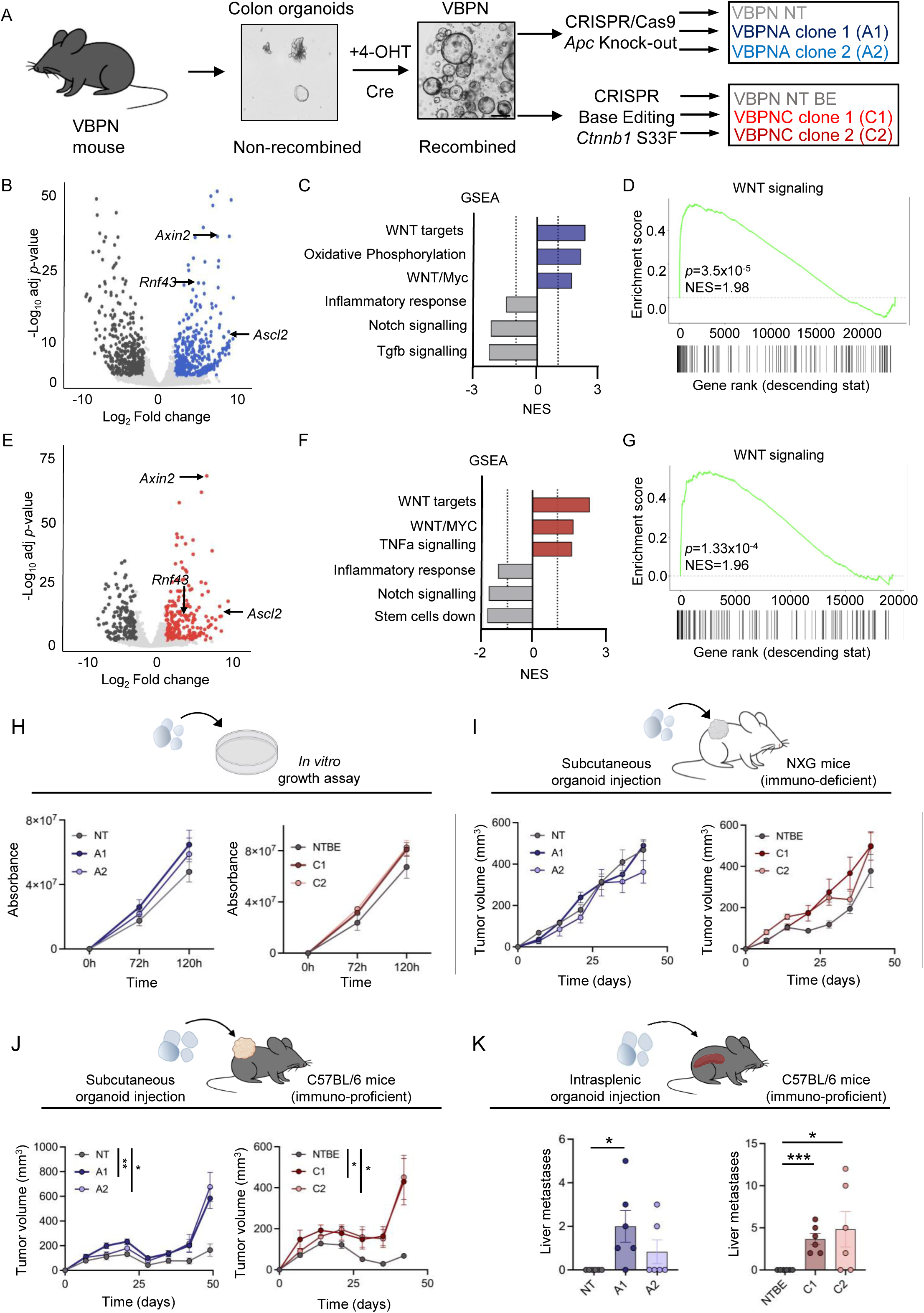
Immune evasion by WNT-activation. **A,** Schematic describing the generation of isogenic VBPNA and VBPNC organoids. Scale bar 250 µm. **B,** Volcano plot of differentially expressed genes from VBPN versus VBPNA organoids. **C,** Gene set-enrichment analysis of VBPN versus VBPNA organoids for indicated gene sets. **D,** Gene set-enrichment analysis of VBPN versus VBPNA organoids for WNT-pathway activation. **E,** Volcano plot of differentially expressed genes from VBPN versus VBPNC organoids. **F,** Gene set-enrichment analysis of VBPN versus VBPNC organoids for indicated gene sets. **G,** Gene set-enrichment analysis of VBPN versus VBPNC organoids for WNT-pathway activation. **H**, Growth assay of indicated organoids *in vitro*. **I,** Tumor growth of subcutaneous tumors in immunodeficient mice (n=6). **J,** Tumor growth of subcutaneous tumors in immuno-proficient mice (n=6). **K,** Number of liver metastases in immuno-proficient mice. Data were analyzed by unpaired t-test. (n=6). Data in H, I and J were analyzed by two-way ANOVA. Adjusted *p* values shown in the enrichment plots in D and G by applying the Benjamini-Hochberg procedure to control the false discovery rate across all tested gene sets.

To phenotypically characterize these organoids, we analyzed their proliferation *in vitro*. Activation of WNT-signaling, did not alter the growth kinetics in comparison to VBPN control organoids (Figure 3H). Next, we tested the response of the organoids to 140 drugs, most of which are FDA-approved (Figure S3A). We also included cells with CRISPR/Cas9-mediated *Rnf43* deletion based on its predictive capacities in patients treated with BRAF/EGFR combinations (27). *Rnf43*-mutant VBPN organoids showed a pronounced response to targeted therapy, while *Ctnnb1*- and *Ape*-mutant VBPN organoids displayed resistance to BRAF and MEK inhibitors such as Cobimetinib, Encorafenib, and Trametinib (Figures S3B-E).

Next, we wanted to follow our observation of immune regulation by WNT-signaling activity. To this end, we transplanted organoids into immunodeficient mice, which revealed no differences in tumor growth depending on WNT-pathway alterations (Figure 3I). However, when transplanted into immuno-proficient C57BL/6 mice, both clones from VBPNA and VBPNC organoids showed significantly increased growth compared to the non-targeting controls (Figure 3J). This phenotype was also observed in a liver colonization assay, with metastases detected only in mice injected with *Ape*-or *Ctnnb1*-mutant VBPN organoids (Figure 3K). These findings suggest that the enhanced WNT-activity leads to immune suppression, enhancing tumor growth and metastasis.

### WNT-activation rewires the tumor immune microenvironment

To characterize the composition of the tumor-ecosystem and its response to WNT-activation, we performed Cellular Indexing of Transcriptomes and Epitopes by Sequencing (CITE-seq) on single-cell level (Figure 4A). We applied a 102-antibody immune panel, enabling comprehensive profiling of immune cell polarization and activation. Single-cell RNA sequencing of 11,559 cells identified five distinct cancer cell clusters alongside major immune cell populations (Figure 4B-C). Profiling of the cancer cell clusters revealed a significant increase of cluster 2 in WNT-activated tumors. Analysis of this cluster demonstrated significant increases in the expression of the WNT-target genes *Axin2* and *Lgr5*. Additionally, WNT-genes signatures were increased in cancer cells found in VBPNA and VBPNC tumors (Figure 4D), which confirms the increased WNT-activation. The immune cell compartment also underwent significant remodeling upon WNT-activation (Figure 4B-C). Neutrophils and macrophages were polarized to pro-tumorigenic phenotypes with myeloid derived suppressor cell and M2 macrophage signatures enriched in WNT-activated tumors (Figure S4A-E). Next, we focused on adaptive immune cells and re-clustered the single cell RNA-seq data to include the surface protein layer of CD4^+^ and CD8^+^ T cells, which led to enhanced resolution of detectable clusters and cell states (Figure 4E). We detected the increase of TCF7⁺ progenitor-like CD4 T cells (Figure 4F), consistent with a stem-like memory phenotype. In addition, we noticed the presence of exhausted CD8 T cells only in WNT-activated tumors, while naive or cytotoxic CD8 T cells were not significantly affected by WNT-alterations (Figure 4F-G). To test the functional role of T cells, we depleted CD4 and CD8 T cells with blocking antibodies after transplantation of VBPNA organoids in C57BL/6 mice (Figure 4H). Tumor growth analysis revealed an anti-tumor phenotype of CD4 T cells (Figure 4I-J, S5A-B). Surprisingly, depletion of CD8 T cells led to reduced tumor growth compared to IgG control (Figure 4I). These findings revealed a contrasting role for T cell subsets, with CD4 T cells limiting tumor progression and CD8 T cells supporting tumor growth. The unexpected pro-tumor effect observed upon CD8 T cell depletion may reflect a predominance of exhausted CD8 T cells within these tumors, which lack cytotoxic function and may contribute to an immunosuppressive microenvironment. In summary, WNT-pathway driven changes in the tumor immune microenvironment led to impaired T cell states that promote tumor progression.

**Figure 4.**
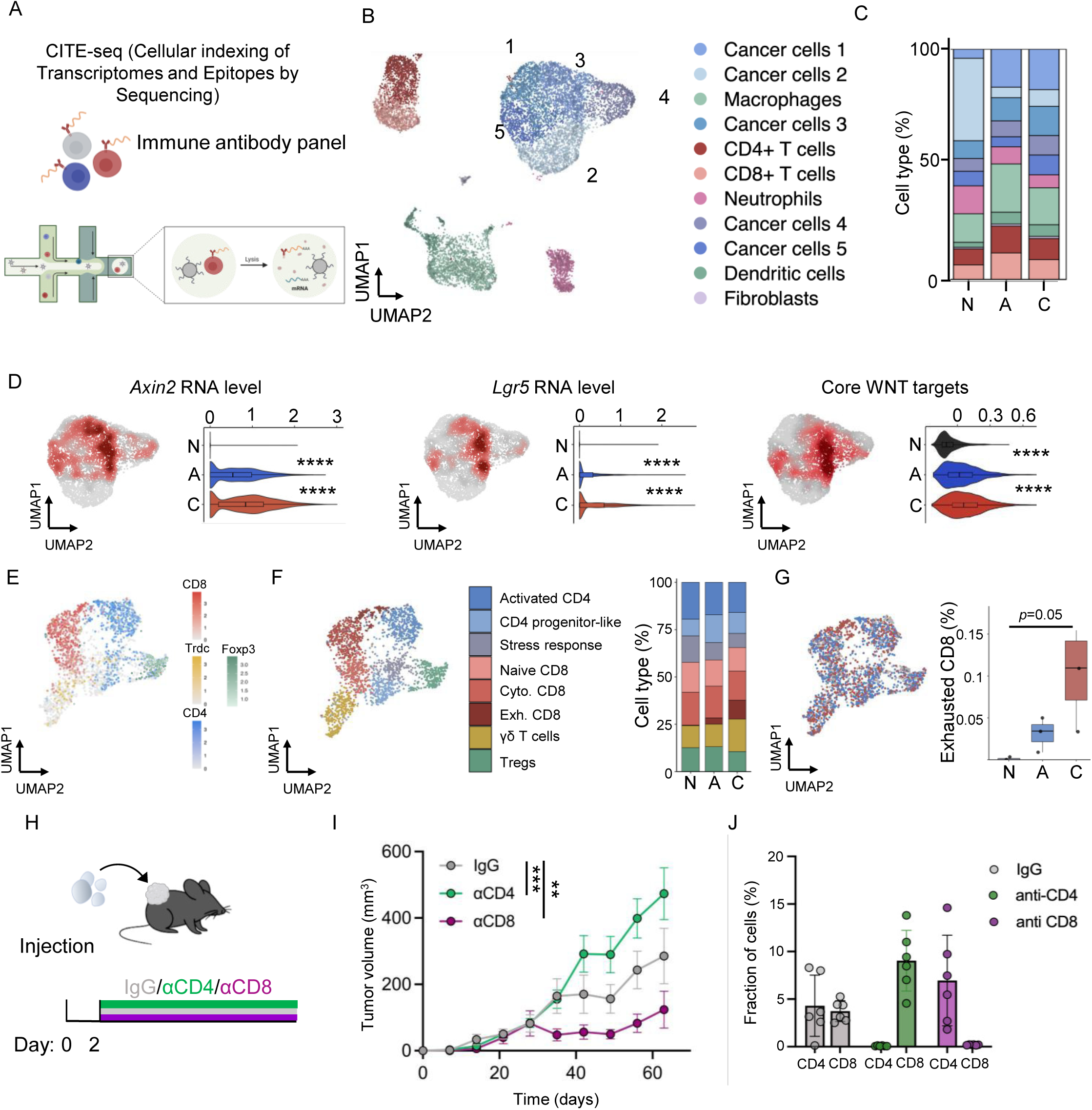
Immune phenotyping of *Brat*-mutant tumors. **A,** Schematic describing the Cellular indexing of Transcriptomes and Epitopes by Sequencing (CITE-seq) workflow. **B,** UMAP plot (11,559 cells) showing the cellular distribution of VBPN, VBPNA and VBPNC tumors. **C,** Bar-plot describing the distribution of cell types across genotypes. N=non-targeting VBPN; A= VBPNA; C=VBPNC. **D,** Quantification of WNT-target gene expression in the epithelial cell compartment of tumors. Statistical significance was assessed by Wilcoxon rank-sum test with Benjamini-Hochberg correction. N=non-targeting VBPN; A= VBPNA; C=VBPNC. **E,** UMAP plot showing T cell annotations. **F,** UMAP and bar-plot showing the quantification of T cell state across genotypes. N=non-targeting VBPN; A= VBPNA; C=VBPNC. **G,** UMAP and box-plot showing CD8 T cell exhaustion cluster across genotypes. Data were analyzed by unpaired *t*-test. N=non-targeting VBPN; A= VBPNA; C=VBPNC. **H,** Schematic describing the experimental set up for T cell depletion *in vivo*. **I,** Tumor growth curve under T cell depletion. Data were analyzed by two-way ANOVA (n=6 per group). **L,** Quantification of T cells depletion at endpoint.

## Discussion

Here we identify a mechanism of WNT-signaling-driven transformation of SSA to serrated CRC. Modeling human-relevant genetic alterations, frequently detected in *BRAF*-mutant serrated CRC, revealed a critical role for Notch1-signaling in driving tumor progression and metastasis to clinically relevant sites such as the liver and lungs. Similar to human serrated CRC, these mouse models acquired WNT-pathway alterations in the course of malignant transformation. WNT-pathway activation was essential for progression by driving immunosuppression.

Our data show that similar to *Kras*-mutant serrated CRC (7), Notch1 activation is crucial to enhance metastatic progression in *Braf*-mutant models. Since we induced the recombination of transgenes with systemic application of tamoxifen, we observed tumors in the entire intestine. These tumors, in contrast to the *Kras*-mutant tumors (7), acquired WNT-pathway mutations. We identified and investigated mutations in genes previously associated to serrated precursors of CRC (10,13,38,39); demonstrating the human relevance of the GEMMs. Our functional characterization of the putative driver genes in the VBPN background showed that *Apc* and *Ctnnb1* mutations led to robust tumor initiation in the intra-colonic injection model. However, deletion of *Rnf43*, despite being associated with serrated CRC, did not lead to tumor initiation, but led to enhanced therapy response, compared to *Apc*- or *Ctnnb1*-mutant organoids. Potential reasons for this might be additional co-occurring genetic events that amplify WNT-signaling in concordance with *Rnf43*-alteration, which are needed for tumor initiation but do not impact therapy responses of established tumors. The observation that *Ape*-deletion is a potent driver of *BRAF*-mutant CRC aligns with previous findings showing that *Apc* alterations define a poor-prognosis subgroup of *BRAF*-mutant CRC (40). However, the association of *Apc* or *Ctnnb1* mutations with a CMS2/3 phenotype suggests that WNT-overactivation leads to better prognosis, of established tumors, compared to VBPN/CMS4 tumors. Collectively, these results provide functional evidence that activation of WNT-signaling contributes to the progression of serrated *BRAF*-mutant CRC.

Our observation that WNT-activation in *BRAF*-mutant CRC suppresses inflammatory and immune-regulatory responses is consistent with correlative analyses. These analyses show that WNT-pathway activation in CRC is associated with immune exclusion (41,42). However, WNT-pathway activation may not be required in all *BRAF*-mutant MSS CRCs, as suggested by our analysis. Evidence from animal models also indicates that serrated CRC can progress without substantial WNT-activation (1). One possible explanation for the WNT-low tumors is that they represent distinct serrated CRC subtypes that evade immunosurveillance through alternative mechanisms, such as the creation of an immunosuppressive, TGF-β-high tumor immune microenvironment, as we have previously shown (7).

Functional investigation of WNT-activation and immune-regulatory properties are also reported in melanoma and hepatocellular carcinoma (HCC) (43,44). In both entities WNT-activation led to failed priming of CD8^+^ T cells, by suppressing dendritic cell recruitment. Interestingly, this recruitment defect was driven by down-regulation of the chemokines CCL4 and CCL5. This indicates that chemokine suppression is a commonly observed mechanism of WNT-induced suppression of anti-tumor immunity. While immune evasion appears to function via a different mechanism in CRC, novel strategies targeting WNT-signaling may still be effective. For example, siRNA encapsulated in lipid nanoparticles targeting *CTNNB1* (LNP-CTNNB1), which induces immune rewiring in HCC (45), could represent a promising approach to inhibit WNT signaling in CRC.

Overall, we show that complex genetic interactions are not merely additive but synergistic, rewiring transcriptional programs. Consequently, altering the tumor microenvironment is critical for tumor cell mediated immune evasion. Understanding this crosstalk and what individual mutations contribute to the cancer cell state and tumor microenvironment is essential for designing rational combination therapies and stratifying patients by functional oncogenic programs rather than single-gene alterations.

## Author contributions

M.M. Validation, investigation, visualization, methodology, writing-review and editing. A.G.M.: Validation, Investigation, Visualization, Methodology, writing-review and editing. J.M.: Validation, Investigation, Methodology. U.P.: Methodology. G.D.: Methodology. N.G.: Methodology. K.S.O.: Methodology. K.E.B.: Methodology. I.G.: Methodology. I.N.: Methodology. A.P.: Methodology. Y.P.: Methodology. M.G.: Methodology. N.G.R. Methodology. J.P.: Methodology. K.S.O.: Methodology. I.C.: Methodology. M.M.: Methodology. J.B.: Methodology. S.O.: Methodology. M.B.; Methodology. R.J.: Validation, investigation, visualization, methodology, funding acquisition, supervision, writing original draft, writing-review and editing.

## Acknowledgment

The authors are grateful for technical support from Carolin Artmann, Gabriele Schmidt and Raja FlOchter. We acknowledge the DKFZ - scOpenLab, Sequencing OpenLab, Genomics Core Facility, flow cytometry facility and Central Animal facilities (ZPF) for excellent service. Fellowships were provided by the Helmholtz International PhD Graduate School to M.M., G.D. and by the Merk’sche Society for Art and Science to A.G.M.. M.M. received a Joachim Herz foundation fellowship. Schematics were created with BioRender.

## Material and Methods

### CRISPR knock-out and base editing vectors

CRISPR genomic engineering vectors were generated via conventional oligo cloning strategies. Plasmids are shown in Table S1. Base Editing (BE) contains a dCas9 domain coupled to a cytosine deaminase (APOBEC). This facilitates the deamination of a cytosine (C), leading to an uracil base (U), which is then read as thymine (T). F2X is a base editor previously designed (46).

### Cell line engineering vectors

Cell lines were generated via lentiviral transduction using the following plasmids shown in Table S2.

### Virus production

Lenti-X™ 293T cells (Takara Bio, #632180) were maintained in DMEM GlutaMax medium (Thermo Fisher Scientific, #31966021) supplemented with 10% fetal bovine serum (Thermo Fisher Scientific, #A5256701) and 1% penicillin-streptomycin (Sigma, #P4458-100ML). Cells were transfected with a polyethylenimine-based protocol (Polysciences, #23966-1), using a 1:1:1 molar ratio of the transfer plasmid, pMD2.G plasmid, and psPAX2 packaging plasmid. Ultracentrifugation of viral supernatants was carried out at 25,000 rpm for 2 hours at 4°C. The lentiviral pellets were stored at -80°C.

### Ethical statement for human samples

The analyses carried out in the study are covered by ethical approval of the Ludwig-Maximilians University Munich (project number: 22-0120) and were performed in accordance with the Declaration of Helsinki.

### MMR/MSI analysis of human samples

MMR/MSI testing was performed as previously described (47). Briefly, immunohistochemical staining for MLH1, MSH2, PMS2 and MSH6 was performed, whereas nuclear loss of expression with retained staining in the non-neoplastic cells as internal control was considered as mismatch-repair deficient (dMMR), corresponding to MSI. In ambiguous cases, confirmatory testing of the five microsatellite markers was performed (BAT25, BAT26, D5S346 [APC locus], D17S250 [p53 locus] and D2S123).

### *BRAF* mutational analysis of human samples

NGS of FFPE tumor tissue was analyzed using multiplex PCR-based AmpliSeq for Illumina panels (Focus Panel and Cancer Hotspot Panel v2; Illumina, San Diego, CA, USA), targeting recurrent hotspot regions and full genes across 50–52 cancer-related genes (48), including *BRAF*.

### Animal experiments

Alleles used in this study are summarized in Table S3. All animal experiments were carried out in accordance with institutional guidelines. The experiments were approved by the regional regulatory body (Regierungsprasidium Karlsruhe, Baden-WOrttemberg, Germany) under the licenses G-148/20, G-235/20, and G-164/22. Mice were maintained in the DKFZ’s SPF animal facilities under standard environmental conditions, including a 12-hour light/dark cycle, controlled temperature (20–24°C), and humidity (45–65%). Animals were housed in individually ventilated cages and had unrestricted access to autoclaved water and a standard rodent diet.

The following genetically engineered mouse models (GEMMs) were generated in this study:

VBP: *Vil1*-creERT2; *Brat*^V600E/+^; *Trp53*^fl/fl^
VBPN: *Vil1*-creERT2; *Brat*^V600E/+^; *Trp53*^fl/fl^; *Rosa26^N1ied^*
VBPNA: *Vil1*-creERT2; *Brat*^V600E/+^; *Trp53*^fl/fl^; *Rosa26^N1ied^, Ape*^fl/+^
VBPNC: *Vil1*-creERT2; *Brat*^V600E/+^; *Trp53*^fl/fl^; *Rosa26^N1icd^*, *Ctnnb1*^fl/+^
VBPNR: *Vil1*-creERT2; *Brat*^V600E/+^; *Trp53*^fl/fl^; *Rosa26^N1ied^*, *Rnt43*^fl/fl^

All strains were maintained on a C57BL6/J genetic background. The *villin1*CreERT2, *Brat*^V600E^, *Rosa26^N1ied^, Ape* and *Ctnnb1* alleles were maintained in heterozygous configuration, while *Trp53,* and *Rnf43* floxed alleles were used in homozygosity.

### Colonoscopy-guided submucosal injection

Intracolonic injection was performed with 4-Hydroxytamoxifen (4-OHT) for VBPN, VBPNA, VBPNC and VBPNR mouse models. Colonoscopic delivery of 4-OHT was performed following established protocols (49). Briefly, mice were anesthetized using isoflurane, and a portable veterinary endoscopy system (Karl Storz, Tele Pack Vet X LED) equipped with a Hopkins telescope (#67030 BA) was used. Injections were administered using a Hamilton syringe (#7656-01), a transfer needle (#7770-02), and a fine injection needle (#7803-05). Three separate injections of 70 µl each were delivered into the colon wall to create visible blebs. Tumor initiation and progression were monitored longitudinally via colonoscopy.

### Generation of mouse organoids

Mouse colon organoids were generated following previously published protocols (49). Briefly, colon tissue was excised, cut into ∼5 mm fragments, and washed repeatedly in cold PBS. After allowing tissue pieces to settle, the supernatant was removed, and the washing step was repeated until the supernatant appeared clear. The fragments were then incubated in PBS containing 25 mM EDTA for colon (Invitrogen, #AM9260G) for 30 minutes on a rotating platform at 4°C. Supernatant fractions were collected and examined, and the cleanest crypt-containing fraction was selected. These were pooled, passed through a cell strainer, and centrifuged to collect the crypts.

The resulting pellet was resuspended in Cultrex BME, reduced growth factor matrix (R&D Systems, #3433-005-01). Colon organoids were cultured in Advanced DMEM/F12 (Gibco, #12634028) supplemented with 1% L-Glutamine (Thermo Fisher Scientific, #25030024), 1% HEPES (Sigma, #H0887-100), 1% penicillin-streptomycin (Sigma, #P4458-100ML), 50% WRN-conditioned medium (prepared as described in (50)), 20% fetal bovine serum (Thermo Fisher Scientific, #A5256701), 50 ng/μl EGF (Peprotech, #AF-100-15), 10 µM Y-27632 (Holzel, #M1817), 0.5 µM A83-01 (Sigma, #SML0788), and 2 µl/ml Primocin (InvivoGen, #ant-pm-2).

### Organoid genome editing

DNA of organoid cultures was edited via CRISPR genetic engineering for the generation of specific knock-outs and base edits. Organoids were cultured to confluency and subjected for plasmid electroporation. The mix was electroporated with NEPA21 electroporator (Nepagene). Electroporation was conducted with poring pulse: 175 V, 5 ms length, 50 ms interval, 2 intervals, 10% decay rate, and positive polarity and transfer pulse: 20 V, 50 ms length, 50 ms interval, 5 intervals, 40% decay rate and +/- polarity.

### Genome editing validation

After formation of organoids from single cells, clonal organoids were picked manually. Genomic DNA was extracted from the organoids using the DNeasy Blood & Tissue Kit (Qiagen, #69506) according to the manufacturer’s protocol. Subsequently the distinct loci were PCR amplified using primers from Table S4 and subjected to Sanger sequencing (Eurofins Genomics). CRISPR-mediated indel frequencies were quantified using the ICE analysis tool provided by Synthego (https://ice.synthego.com/). Due to the clonality of the cultures, indel frequencies of 0% (wild type, no editing), 50% (editing on one allele) or 100% (editing on both alleles) were expected. Organoid cultures with an editing efficiency of 100% (full knock-out/edit) were taken for further experiments.

### *In vitro* organoid growth

To measure the proliferation rates of different organoids, CellTiter Blue experiments were conducted. Three wells for each organoid line were incubated for 3 hours with CellTiter Blue solution from Promega (#G8081) at a dilution of 1:6 with complete Advanced DMEM media. Following the 3 hour incubation, 100 μl of the supernatant were collected from each well. The UV emission of this plate was then recorded in the spectrometer plate reader, with the settings: 560 nm excitation and 590 nm absorbance. The growth curve was generated using the absorbance values at the indicated time points.

### Subcutaneous injection, tumor growth assay and T cell depletion

Dissociated organoids were resuspended in 100 µl of a 1:1 mixture of PBS and BME. Next, mice were anesthetized and the single-cell organoid suspension was injected into the flank. CD4/8 T cell depletion was performed by treating the mice with 100 µg anti-CD4 antibody (BioXCell, BXC-BE0003-1), anti-CD8alpha antibody (BioXCell, BXC-BE0061) or rat IgG2b isotype control (BioXCell, BXC-BE0061) twice per week via intraperitoneal injections. Treatment started 2 days after organoid injection.

### Intra-splenic injection

Dissociated tumor cells in PBS were slowly introduced into the spleen using a fine-gauge syringe. Access to the spleen was achieved through a small surgical incision in the left lateral abdomen, followed by gentle exteriorization of the spleen. Animals were euthanized four weeks after implantation of the modified tumor organoids.

### Flow cytometry

Organoids were dissociated into single cells using previously described methods. The isolated cells were washed with cold PBS and subsequently resuspended in FACS Buffer. For VBPN organoid lines expressing the TOP-GFP.mc reporter, flow cytometry analysis was conducted immediately following the dissociation into single cells. Analysis were performed on the BD LSR Fortessa flow cytometer (BD Biosciences) and FlowJo 10.8.1 software.

### Immunofluorescence

Following tissue processing, antigen retrieval was performed by steaming the slides in boiling Target Retrieval Solution pH9 (S2367) for 30 minutes. Afterwards, the slides were cooled for 20 minutes at room temperature, rinsed in distilled water and washed in PBST 0.1% for 5’. Next, sections were blocked in TNB buffer (0.1 M Tris-HCl, pH 7.5, and 0.15 M NaCl containing 0.5% w/v blocking reagent, Perkin Elmer #FP1020) for 1–2 hours at room temperature. For immunostaining, slides were incubated for 1 hour at room temperature or overnight at 4°C with primary antibodies (Table S5). Following incubation, sections were washed in PBST 0.1% and then incubated with the appropriate secondary antibody (Table S5) for 30 minutes at room temperature, washed with PBST 0.1% and incubated with DAPI (1:1000) for 10 minutes at room temperature. The slides were washed again in PBST 0.1% and mounted with Fluoromount-G (Southern Biotech #0100-01). Images were acquired using a Zeiss LSM 710 microscope and analyzed with FIJI (ImageJ) or QuPath-0.4.3.

### Immunohistochemistry

Following tissue processing, antigen retrieval was performed by steaming the slides in boiling Target Retrieval Solution pH9 (S2367) for 30 minutes. Following initial tissue processing, antigen retrieval was performed by placing the slides in a steamer containing boiling Target Retrieval Solution (Dako, #S169984) for 30 minutes. The slides were then cooled for 20’ at room temperature, washed in TBST 0.1% and subsequently incubated in 3% hydrogen peroxide (Sigma, #95321) for 10’. Tissue sections were then incubated in TNB Normal Swine Serum blocking buffer, composed of 0.1 M Tris-HCl (pH 7.5), 0.15 M NaCl, and 0.5% w/v blocking reagent (Perkin Elmer, #FP1020Biozol, #LIN-ENP9010-10), at room temperature for 1 hour. Primary antibodies (Table S5) were applied either overnight at 4°C or for 1 hour 90 minutes at room temperature. After incubation, slides were rinsed two times with TBS-T 0.1% and incubated for 30 minutes to one 1 hour at room temperature together with the appropriate biotin-conjugated secondary antibody (Table S6). Signal detection was conducted Then, the slides were washed with TBST 0.1% and incubated with the VECTASTAIN® ABC kit (Vector Laboratories, #PK-6100) for 30 minutes at room temperature., The slides were washed again with TBST 0.1%. following the manufacturer’s instructions. Signal detection was conducted with DAB chromogen (Agilent Dako, #GV82511-2), following the manufacturer’s instructions.

### *In situ* hybridization

*In situ* hybridization (*ISH*) was performed using the manual RNAscope 2.5 HD detection kit (Advanced Cell Diagnostics, Hayward, CA, #32250 RNAscope® 2.5 HD Detection Reagents-BROWN (Advanced Cell Diagnostics, Hayward, CA, #322300) strictly according to the manufacturer’s instructions. Staining was performed on 4 μm formalin fixed paraffin sections, which were cut and then placed in a 60°C oven for 2 hours prior to staining. To ensure the quality and integrity of the available RNA the tissue being investigated was tested with the positive control probe (PPIB). Evaluation was performed only after positive probe quality control results were obtained. To further ensure accuracy and integrity of the staining a negative control probe (DapB, Advanced Cell Diagnostics, Hayward, CA, #0043) was used to confirm that the tissue staining seen was accurate due to binding with the target probe and not non-specific. Probes: *Lgr5* (Advanced Cell Diagnostics, Hayward, CA, #312171), positive control probe *Ppib* (Advanced Cell Diagnostics, Hayward, CA, #321651).

### Tumor dissociation

To obtain single-cell suspensions from tumor tissue, frozen tumor fragments were thawed and subjected to enzymatic digestion using the Tumor Dissociation Kit (Miltenyi Biotec, #130-095-929) according to the manufacturer’s protocol. Briefly, 5 ml of DMEM GlutaMax (Thermo Fisher Scientific, #31966021) together with 500 µl of the enzyme mix was transferred into a gentleMACS C Tube (Miltenyi Biotec, #130-096-334). Tumor pieces were dissociated in C Tubes on the gentleMACS Octo Dissociator with heaters (Miltenyi Biotec, #130-096-427). Processing was performed using the pre-set dissociation program "37C_m_TDK_1" to ensure efficient mechanical and enzymatic breakdown of the tissue.

### Quantitative PCR

RNA was extracted from organoid cultures or other cell populations using the RNeasy Mini Kit (Qiagen, #74106) following the respective manufacturer’s protocols.

cDNA was synthesized from up to 1 μg of total RNA using either the High-Capacity cDNA Reverse Transcription Kit (Applied Biosystems, #4374966) following the supplier’s guidelines. Thermal cycling for reverse transcription involved incubation at 25 °C for 10 minutes, extension at 37-42 °C for 60-120 minutes, and enzyme inactivation at 85 °C.

qPCR reactions were set up using the PowerUp™ SYBR™ Green Master Mix (Thermo Fisher Scientific, #A25778). Primers are listed in Table S6. Reactions were run in technical triplicates on a QuantStudio™ 5 Real-Time PCR System (Applied Biosystems). Thermocycling included initial UDG activation, denaturation at 95 °C, and 40 cycles of annealing/extension, followed by melt curve analysis to confirm product specificity.

Gene expression was normalized to housekeeping genes such as s18 or Gapdh for murine samples. Relative quantification was calculated using the ΔΔCt method, and fold changes were expressed relative to the indicated control conditions.

### CITE-seq analysis

Tumor-derived single-cell suspensions were prepared as previously described and sequencing sample preparation was conducted as in 10X single-cell RNA sequencing. Minimum of 500,000 cells were subjected to further processing for CITE-seq. The TotalSeq™-B Mouse Universal Cocktail, V1.0 (Biolegend, #199902) was prepared according to the manufacturer’s protocol and the cells were incubated with the antibody cocktail for 30 minutes on ice. Afterwards, the cells were washed three times with FACS buffer and resuspended in 100 µl PBS, 2% FBS. CITE-seq libraries were prepared using the Chromium Next GEM Single Cell 3’ Reagent Kit v3.1 (Dual Index), generating one library for gene expression (GEX) and one for cell surface protein together with hashtag oligo (CITE + HTO) detection per sample. CITE-seq data were processed in parallel with gene expression (GEX) and hashtag oligo (HTO) data using the Seurat framework. Antibody-derived tag (ADT) counts generated from TotalSeq™-B antibodies were loaded using the Read10X function and stored as a separate assay within each Seurat object. ADT data were normalized using centered log-ratio (CLR) transformation across cells with Seurat’s NormalizeData function (normalization.method = "CLR"). Following quality control and HTO-based sample demultiplexing, a multi-modal integration was performed using Seurat’s Weighted Nearest Neighbor (WNN) algorithm (FindMultiModalNeighbors), combining transcriptomic and surface protein information for improved clustering and dimensionality reduction. The integration of RNA, ADT, and HTO modalities enabled robust assessment of cellular subtypes and improved resolution of subtle phenotypic differences driven by genetic background. Single-cell libraries were sequenced on an Illumina NovaSeq 6000 platform using a paired-end, dual-indexing strategy. Sequencing was performed with the following cycle configuration: 28 cycles for Read 1, 10 cycles each for the i7 and i5 index reads, and 90 cycles for Read 2. The target sequencing depth was approximately 30,000 read pairs per cell for gene expression libraries and around 500 read pairs per cell for each hashing and CITE-seq antibody.

### RNA sequencing

RNA from tumor and organoid samples was conducted with RNeasy Mini RNA Isolation Kit according to the manufacturer’s guidelines (Qiagen, #74106). The Bioanalyzer (Agilent) and the Bioanalyzer RNA Kit (Agilent, #5067-1513) were used to assess RNA quality and amount. Whole transcriptome library preparation was conducted with an adapted protocol based on SMART-seq2. Reverse transcription was performed with 1 µM template switching oligo (IDT), 1 mM dithiothreitol (Takara Bio Clontech, # 707265ML), 1 U/µl RNase Inhibitor (Promega, #N2615), Maxima H Minus Reverse Transcriptase (LIFE Technologies, #EP0753) and 1x Maxima H Minus Reverse Transcriptase buffer. Afterwards, tagmentations of cDNA and further library preparation was conducted with Nextera XT DNA Library Preparation Kit (Illumina, FC-#121-1030).

Raw sequencing reads were aligned to the appropriate reference genome (mouse genome: refdata-gex-mm10-2020-A) using STAR (v2.7.9a) with default parameters. Read quality and adapter contamination were assessed using FastQC (v0.11.9) and aggregated with MultiQC (v1.13). Aligned reads were quantified using featureCounts (subread package, v2.0.3), and gene-level count matrices were generated based on Ensembl gene annotations.

Subsequent analyses were performed in R (v4.3.1). Low expressed genes were filtered out using a minimum count threshold across samples. Data normalization, variance stabilization and differential expression analysis were carried out using the DESeq2 package (v1.40.2). Principal component analysis (PCA) and sample clustering were used to assess global transcriptomic variation. Significantly differentially expressed genes were defined using a Benjamini-Hochberg adjusted p-value < 0.05 and absolute log₂ fold-change > 1, unless stated otherwise. Gene Ontology (GO) and pathway enrichment analyses were performed using fgsea and visualizations (e.g. volcano plots, heatmaps) were generated with ggplot2 and pheatmap.

RNA-seq libraries were pooled and sequenced with Illumina NextSeq and a mid-output flow cell with single-end 75 bp read sequencing. A total of 1.4 pM with 1% PhiX was loaded on the flow cell.

### Tumor bulk CMS classification

Bulk RNA-seq count matrices were used to assign CMS labels. Mouse gene symbols were first mapped to human orthologs and/or Entrez identifiers to match the required CMS reference gene space; duplicated and unmapped identifiers were removed prior to classification. CMS calling was then performed using the MmCMS/CMScaller framework (template-based nearest template prediction with an FDR threshold of 0.05), yielding per-sample CMS1–CMS4 predictions and corresponding subtype similarity scores. To visualize subtype distributions across tumor cohorts (VBPN, VBPNA, VBPNC), CMS calls were summarized as proportions and displayed as stacked bar plots. In addition, CMS similarity score matrices (CMS1–CMS4) were plotted as heatmaps (column-scaled where indicated), with samples annotated by experimental group to facilitate comparison of CMS program strength across conditions.

### Whole exome sequencing

Whole exome sequencing libraries of tumors were generated using the SureSelectXT Automation Reagent Kit following the manufacturer’s instructions. In brief, 200 ng of gDNA was fragmented to ∼150bp using a Covaris LE220 ultrasonicator (Covaris, Inc.). Subsequently, library preparation was performed on a Bravo automated liquid handler (Agilent Technologies) including end-repair, A-tailing, adaptor ligation and amplification. The concentration of amplified, adaptor-ligated DNA library was determined using the TapeStation (Agilent Technologies). In the subsequent steps 750 ng of amplified, adaptor-ligated DNA library was used for the hybridization reaction with the SureSelectXT All Exon v7 bait set. The DNA-library/bait hybrids were captured using streptavidin-coated magnetic beads (Dynabeads MyOne Streptavidin T1 by Thermo Fisher Scientific). Index tags were added in the course of PCR-amplification of the captured libraries. The final libraries were validated using Qubit (Invitrogen) and Tapetstation (Agilent Technologies). 2x 100 bp paired-end sequencing was performed on the Illumina NovaSeq 6000 according to the manufacturer’s protocol.

The resulting datasets were aligned and called for SNVs and indels as described before (51). In brief, alignment was performed by BWA MEM. SNVs and indels were called using Strelka2, mouse tumors were paired with normal tissue from the same donor for somatic mutation calling. Variants and predicted effects were annotated using SnpEff. Variant call format (VCF) files were filtered to retain only high-confidence variants (“PASS” filter). Variant allele frequencies were calculated and variants in the genes as described to (52) were extracted.

### Mutational profiling of *BRAF*-mutant CRC from publicly available data

Somatic mutation data were obtained from publicly available datasets via the cBioPortal for Cancer Genomics (https://www.cbioportal.org). We downloaded the Colorectal Adenocarcinoma (TCGA, PanCancer Atlas) (34) and Colorectal Cancer (MSK, Gastroenterology 2020) (35) datasets. Data were subset to include only *BRAF*-mutant, microsatellite-stable (MSS) tumors. Somatic mutations in key driver genes (*APC*, *BRAF*, *TP53*, *CTNNB1*, *RNF43*, *AXIN2*) were extracted, and an oncoplot summarizing mutation frequency and distribution across samples was generated using the maftools R package (https://bioconductor.org/packages/release/bioc/html/maftools.html) (35))

### Drug screening of murine cancer organoids

Drug screening of the murine organoids was performed as previously described (53). The drug library was composed of 140 drugs, most of which are FDA-approved for clinical use. Each compound was tested at five concentrations in a 5-fold dilution series. The drugs were randomly arranged in two 384-well screening plates. Control wells were distributed across both library screening plates and treated with either DMSO (0.1%) or Bortezomib (5 µM) as negative or positive control, respectively. All drugs were obtained from Selleck Chemicals.

Drug treatment was performed 3 days after seeding by first aspirating medium and then adding 45 µl of fresh medium and 5 µl of previously diluted drug solution using a Biomek FX robotic device (Beckman Coulter).

Quantification of cell viability following 96 hours of drug treatment was performed using the CellTiter-Glo® Luminescent Cell Viability Assay (Promega, #G9682). Again, medium was discarded and 30 μl of CellTiter-Glo solution was added to each well. The plates were incubated at room temperature for 30 minutes. Luminescence was measured using a Mithras LB 940 microplate reader.

### Analysis of drug sensitivities

Organoid drug sensitivity was calculated by normalizing all luminescence values to the DMSO wells from the respective experiment. Area under the curve (AUC) values were calculated for each organoid line and drug by plotting normalized cell viability against drug concentration. Drug concentrations were normalized to range from 1 (maximum concentration) to 0.2 (highest dilution) and trapezoid integration was performed using the R package pracma (https://cran.r-project.org/web/packages/pracma/index.html).

### Statistical analysis

Statistical analysis of single-cell RNA sequencing (scRNA-seq) data was performed using R (version 4.3.3) and the Seurat package (version 5.0.2), as outlined in the respective analysis sections. For comparisons of gene expression or gene set activity between cell groups (e.g., split violin plots), the Wilcoxon rank-sum test was applied as implemented in Seurat. *P*-values were adjusted for multiple comparisons using the Benjamini-Hochberg method unless otherwise specified. Additional statistical analyses were conducted in GraphPad Prism version 9 (GraphPad Software), unless stated otherwise. The statistical tests used, number of biological replicates, and sample sizes are provided in the corresponding figure legends. Data are presented as mean ± standard error of the mean (SEM). Statistical significance is indicated as follows: \**p* < 0.05, \*\**p* < 0.01, \*\*\**p* < 0.001, and \*\*\*\**p* < 0.0001.

---

**Figure S1.**
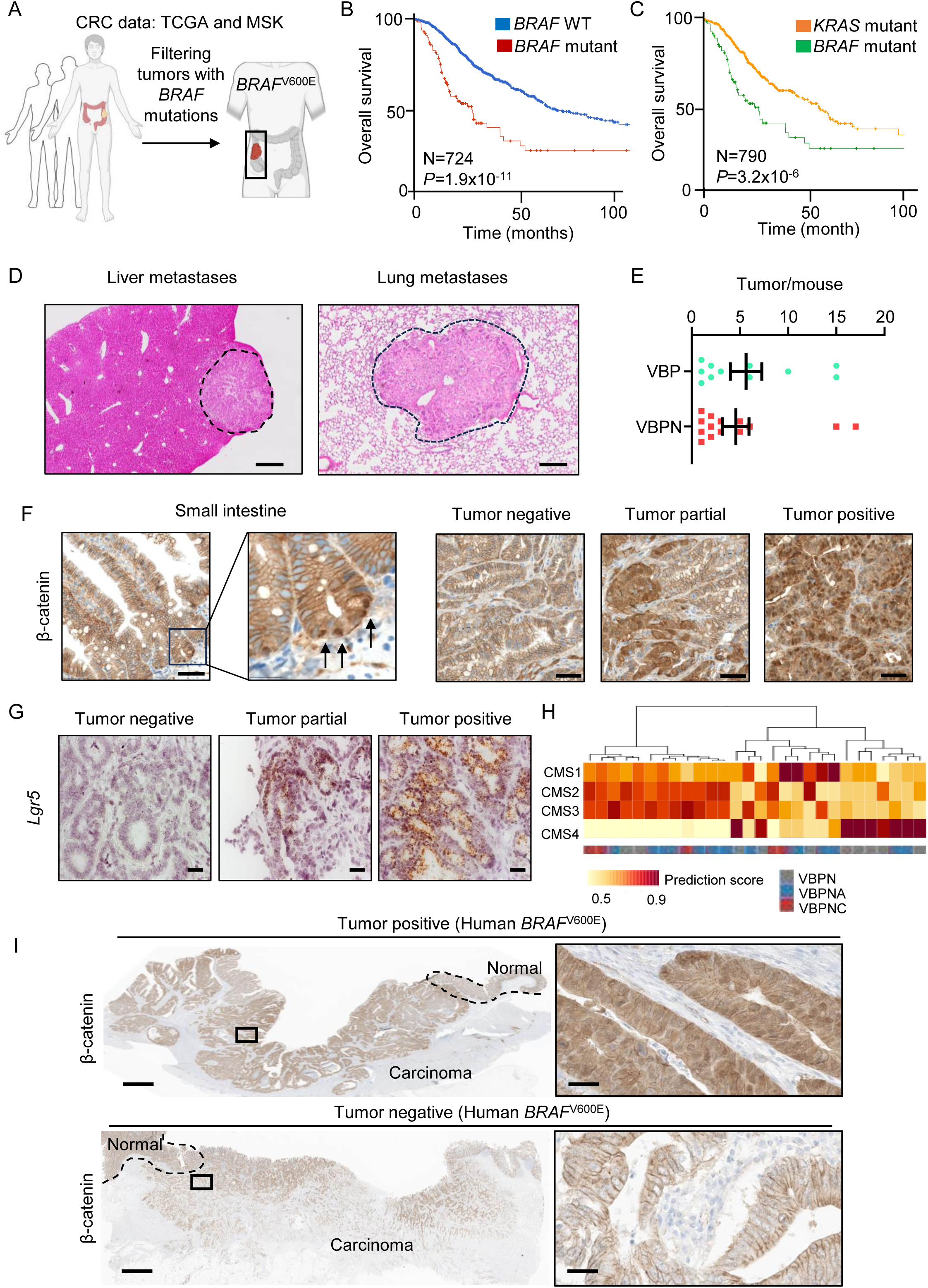
Phenotypic and molecular features of *Braf*-mutant tumors. **A,** Schematic describing the *BRAF*^V600E^-mutant cohort from TCGA and MSK-datasets. **B,** Kaplan-Meier curve describing survival of *BRAF*-mutant versus *BRAF*-WT. Statistical significance was assessed by two-sided log-rank (Mantel-Cox) test. **C,** Kaplan-Meier curve describing survival of *BRAF*-mutant versus *KRAS*-mutant. Statistical significance was assessed by two-sided log-rank (Mantel-Cox) test. **D,** Representative H&E images of liver and lung metastases from VBPN mice. Scale bar 100 µm. **E,** Quantification of tumors per mouse (VBP n=11; VBPN n=14). **F,** Representative pictures of β-catenin IHC in small intestine and VBPN tumors. Scale bar 50 µm. **G,** *In situ* hybridization pictures of *Lgr5* in VBPN tumors. Scale bar 50 µm. **H,** Heatmap showing the model specific clustering for consensus molecular subtypes. **I,** Representative pictures of β-catenin IHC in human *BRAF*-mutant MSS tumors. Scale bars: left 2 mm, right 20 µm.

**Figure S2.**
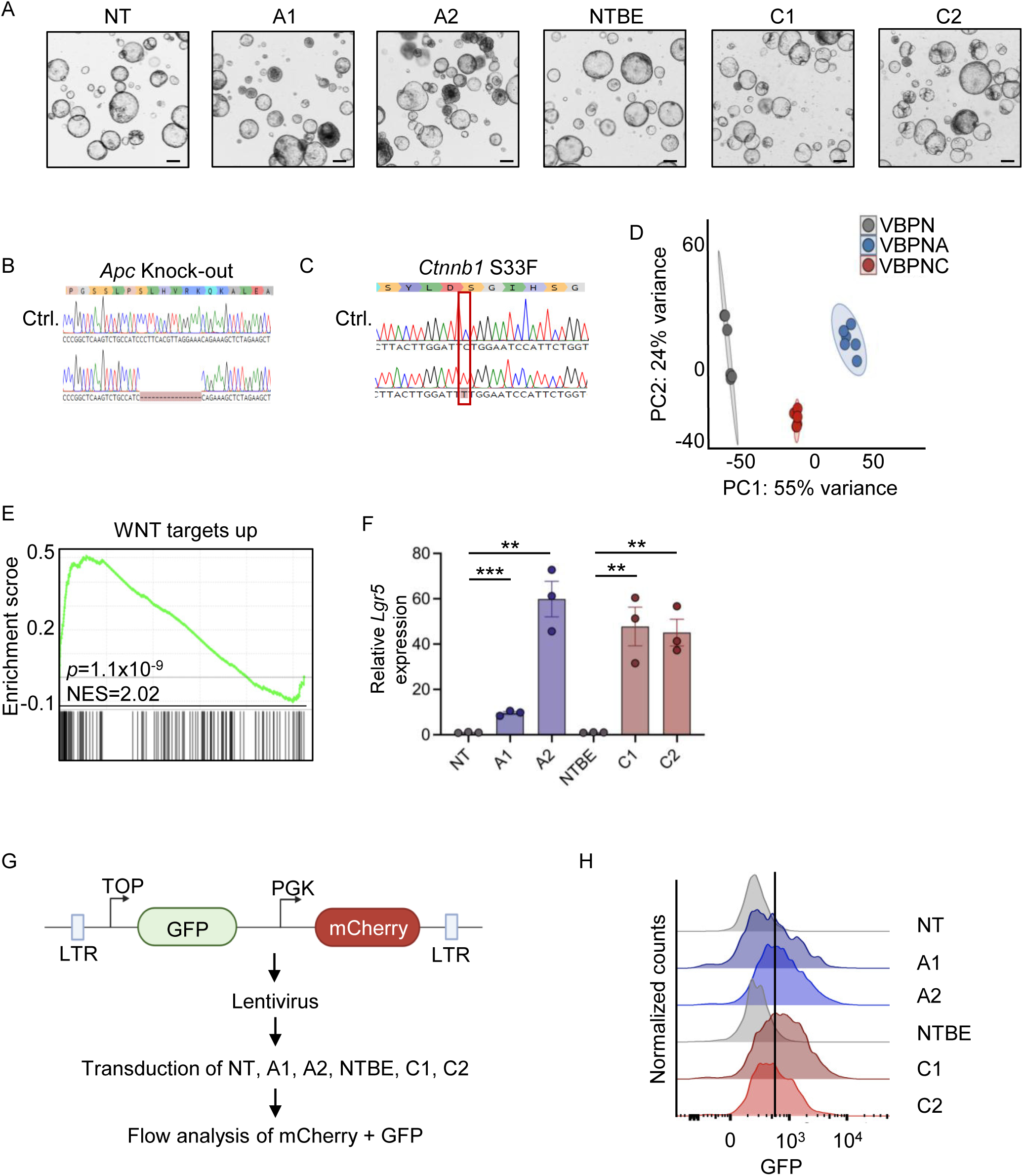
Genome editing and characterization of *Braf*-mutant organoids. **A,** Light microscopy pictures of organoids with indicated genotypes. Non-targeting=NT; Non-targeting base editor=NTBE. Scale bar 200 µm. **B,** Sanger sequencing chromatograms of *Ape* mutation in VBPNA organoids. **C,** Sanger sequencing chromatograms of *Ctnnb1* mutation in VBPNC organoids. **D,** Principle component analysis (PCA) of RNA-seq from organoids. **E**, Gene set-enrichment analysis of VBPN versus VBPNA/VBPNC organoids for WNT-pathway activity. **F,** qPCR analysis of WNT-target *Lgr5* across indicated genotypes. Data were analyzed by unpaired t-test (n=3). **G,** Schematic describing the generation of TOP-GFP WNT-reporter organoids. **H,** Histograms from flow-cytometric analysis of GFP across organoids.

**Figure S3.**
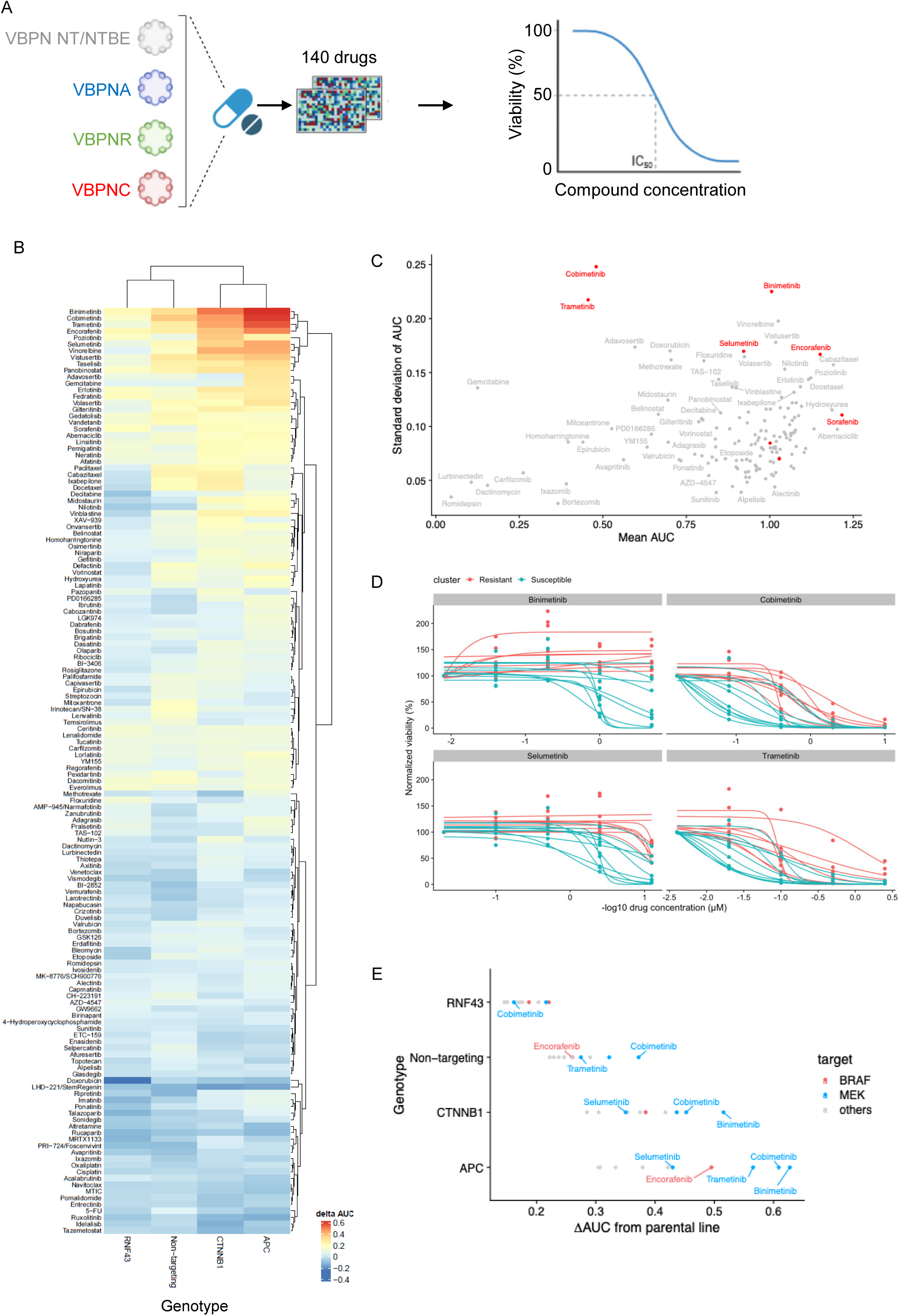
Drug screen in *Braf*-mutant organoids. **A,** Overview of the drug screening experiment of murine organoid lines and a library of 140 drugs. Each organoid line was screened in duplicates. **B,** Heatmap displaying the mean delta AUC values for all drugs. Per genotype, the mean AUC values per drug of two independent clones were calculated and subtracted from the mean AUC values of the parental organoid line. **C,** Mean AUC of all tested drugs plotted against the standard deviation of AUC, BRAF and MEK inhibitors are highlighted. **D,** Dose-response curves for selected MEK and BRAF inhibitors colored by resistant (*Ape*, *Ctnnb1*) and susceptible (*Rnf43*, Parental, VPBN-NT, VBPN-NTBE) genotypes. **E,** Difference in AUC (ΔAUC) of each genotype compared to the parental organoid line.

**Figure S4.**
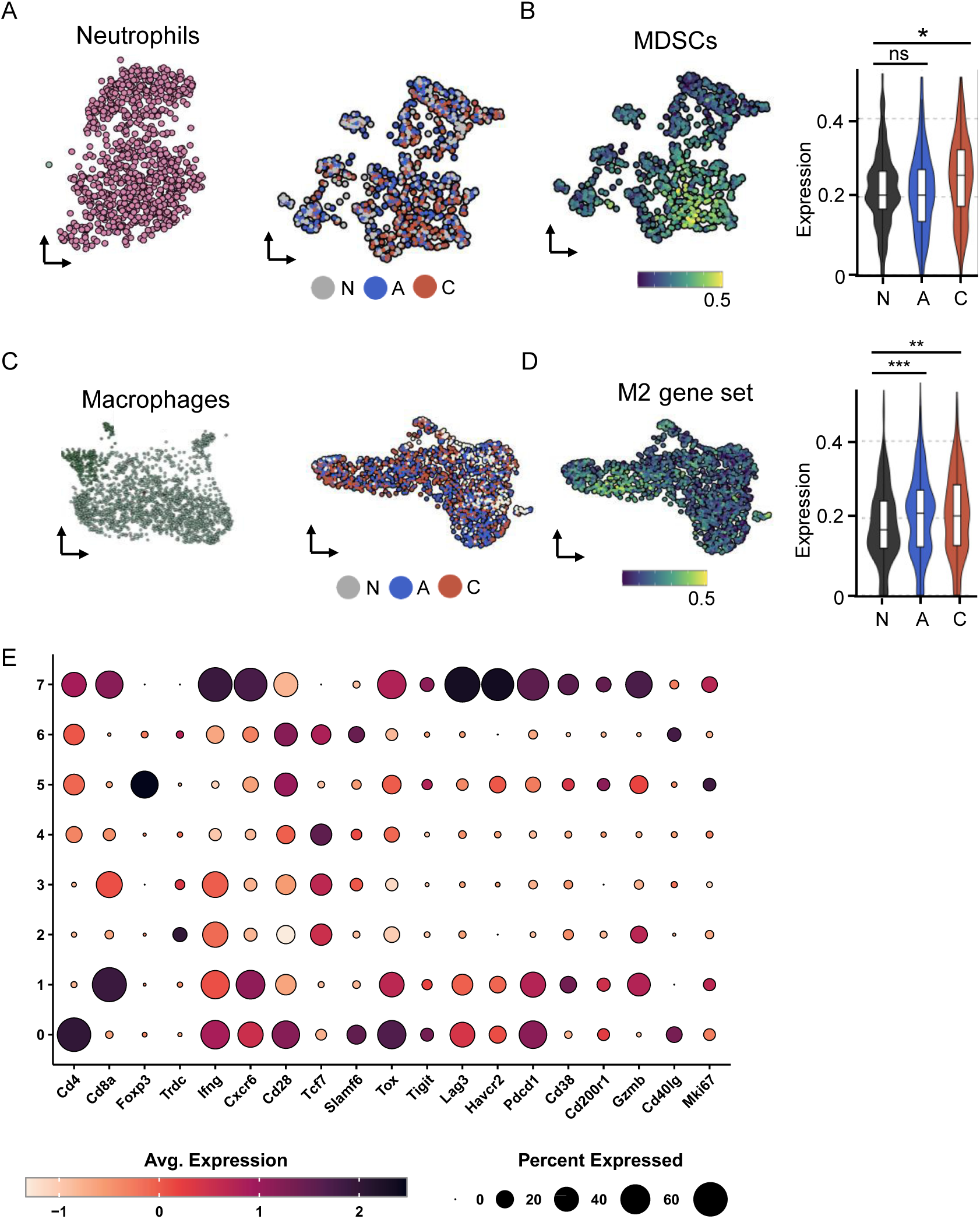
CITE-seq characterization. **A,** UMAP plot showing neutrophil population across genotypes. **B,** Quantification of myeloid derived suppressor cell signatures across genotypes. Statistical significance was assessed by Wilcoxon rank-sum test with Benjamini-Hochberg correction. **C,** UMAP plot showing macrophage population across genotypes. **D,** Quantification of M2 macrophage signatures across genotypes. Statistical significance was assessed by Wilcoxon rank-sum test with Benjamini-Hochberg correction. **E**, Bubble plot showing cell type annotations of the CITE-seq analysis. In A, B, C and D N=non-targeting VBPN; A= VBPNA; C=VBPNC.

**Figure S5.**
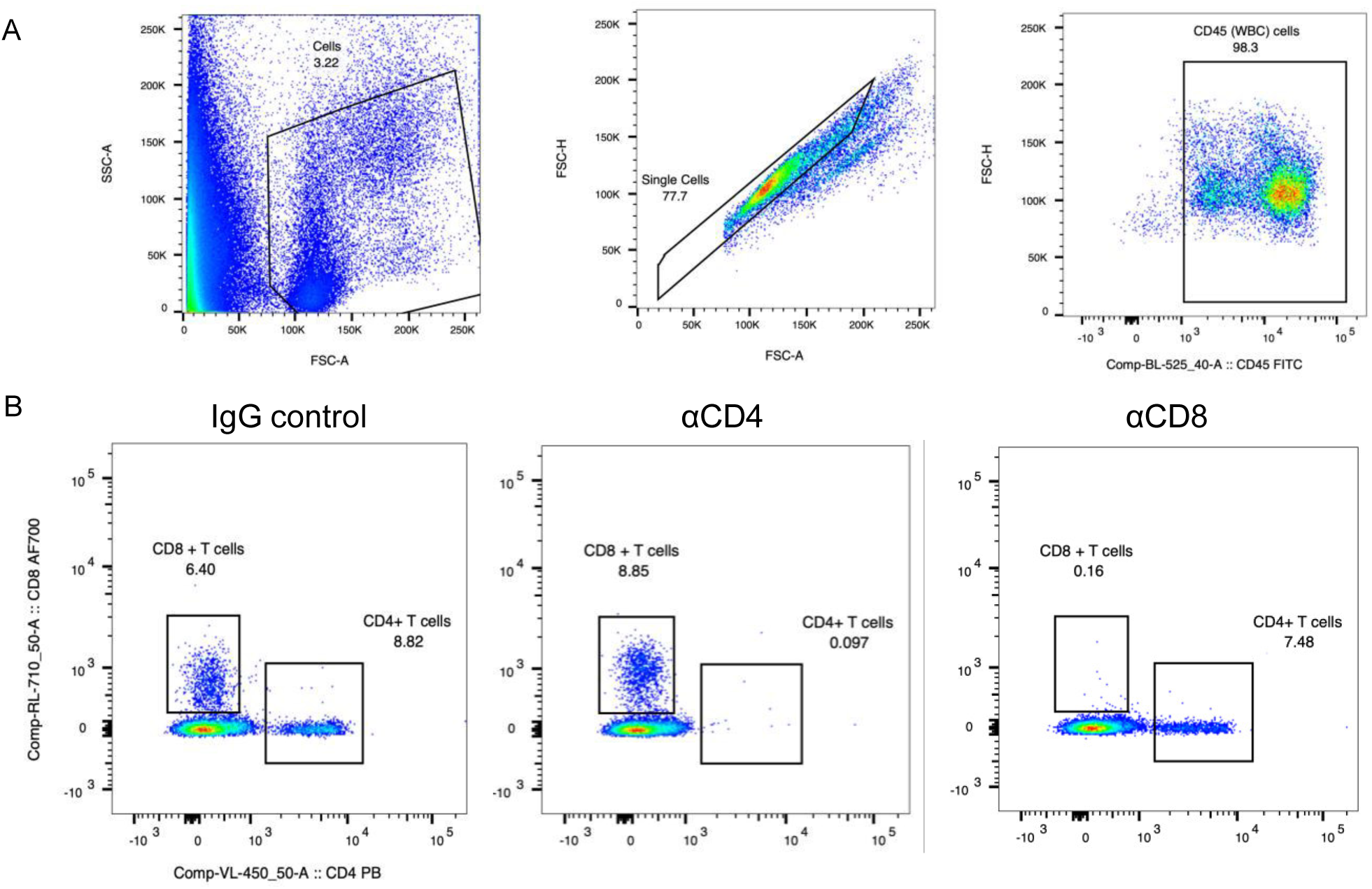
T cell depletion. **A,** Flow cytometric analysis and gating strategy for T cell depletion analysis. **B**, Representative plot showing CD4 or CD8 T cell depletion.

